# Tracking mitochondrial density and positioning along a growing neuronal process in individual *C. elegans* neuron using a long-term growth and imaging microfluidic device

**DOI:** 10.1101/2020.07.27.222372

**Authors:** Sudip Mondal, Jyoti Dubey, Anjali Awasthi, Guruprasad Reddy Sure, Sandhya P. Koushika

## Abstract

The long cellular architecture of neurons requires regulation in part through transport and anchoring events to distribute intracellular organelles. During development, cellular and sub-cellular events such as organelle additions and their recruitment at specific sites on the growing axons occur over different time scales and often show inter-animal variability thus making it difficult to identify specific phenomena in population averages. To measure the variability in sub-cellular events such as organelle positions, we developed a microfluidic device to feed and immobilize *C. elegans* for high-resolution imaging over several days. The microfluidic device enabled long-term imaging of individual animals and allowed us to investigate organelle density using mitochondria as a testbed in a growing neuronal process *in vivo*. Sub-cellular imaging of an individual neuron in multiple animals, over 36 hours in our microfluidic device, shows the addition of new mitochondria along the neuronal process and an increase in the accumulation of synaptic vesicles at synapses, both organelles with important roles in neurons. Long-term imaging of individual *C. elegans* touch receptor neurons identifies addition of new mitochondria and interacts with other moving mitochondria only through fission and fusion events. The addition of new mitochondria takes place along the entire neuronal process length and the threshold for the addition of a new mitochondrion is when the average separation between the two pre-existing mitochondria exceeds 24 micrometers.

## Introduction

Development and maturation of neurons occur through the processes of cell birth, cell differentiation, cell migration, neuronal process outgrowth, dendritic arborization, synaptic growth, and organelle transport (Smith et al., 2010, Hobert, 2010, Varier and Kaiser, 2011, Smith and Gallo, 2017). Complex cellular architectures present in polarized cells such as neurons need transport of organelles to support many energy-dependent processes such as neurotransmission. Neurotransmission heavily relies on regulated transport of organelles to meet metabolic demand (Sheng and Cai, 2012). One such organelle is mitochondria that are essential for neuronal function and thought to contribute to various disease pathologies (Liu et al., 2012, Millecamps and Julien, 2013). Of the large number of mitochondria in a neuronal process, only ∼20% move and the halting events depend on molecular motors, calcium, and other signals that are read by the molecules regulating mitochondrial transport (Chen and Sheng, 2013, Kang et al., 2008, Sun et al., 2013, Schwarz, 2013, Misgeld and Schwarz, 2017). Mitochondrial trafficking is a dynamic process that continues throughout the development when several neuronal processes undergo several folds elongation during the growth of the animal. Thus, mitochondria offer a unique opportunity to demonstrate the utility of long-term imaging in individual neurons. Thus far, invertebrate axons typically have not been reported to have any general anatomical features such as nodes of Ranvier with vast numbers of mitochondria present in vertebrate neurons (Ohno et al., 2011, Chiu, 2011). Multiple neurons, both *in vivo* and in non-myelinated neurons in culture, show that the density of mitochondria is invariant (Misgeld and Schwarz, 2017, Wang and Schwarz, 2009a, Morsci et al., 2016, Sure et al., 2018). Majority of mitochondria in *C. elegans* touch neurons have been reported to be present at actin-rich regions that are distributed throughout the neuron (Sood et al., 2018); however, the density of mitochondria in adult animals remains largely invariant (Sure et al., 2018, Morsci et al., 2016). Despite averages being tightly regulated, inter-animal variability does exist. For example, notwithstanding nearly invariant cell lineages variability in cell-cycle onset and duration of *C. elegans* vulval precursor cells have been observed (Keil et al., 2016). Resolving cellular and sub-cellular events at time scales ranging from minutes to several days allow for a deeper understanding of several processes like development and aging that show variability in mitochondrial density and axonal length in a clonal genetic population (O’Toole et al., 2008), heterogeneity of mitochondrial dynamics within individual cells (Sison et al., 2017), and differential mitochondrial dynamics and distribution within different segments of a neuron (Obashi and Okabe, 2013). To investigate where mitochondria are added one needs to image the same animal over multiple days and analyze the specific location of each mitochondrion over and above the natural variability in individual mitochondrial positions seen during the development.

Developmental events can be easily tracked using *C. elegans*, a well-characterized transparent genetic model, small in size, completely annotated nervous system, and easy to grow in liquid media. These advantages make it an excellent model system to use in microfluidic platforms. Several microfluidic platforms have been developed for *C. elegans* to carry out high-resolution imaging (Chalasani et al., 2007, Caceres Ide et al., 2012, Mondal et al., 2011, Mondal et al., 2012), high-throughput screening (Chung et al., 2008, Mondal et al., 2016, Mondal et al., 2018), long-term imaging (Hulme et al., 2010, Krajniak and Lu, 2010, Xian et al., 2013, Keil et al., 2016, Atakan et al., 2019), and neuronal axotomy (Guo et al., 2008, Samara et al., 2010). Microfluidic device immobilization avoids alterations in *C. elegans* physiological processes compared to anesthetic immobilization (Guo et al., 2008, Mondal et al., 2011, Mondal et al., 2012, Mondal and Koushika, 2014). *C. elegans* can be cultured throughout its entire life cycle by feeding them with bacterial cultures inside microfluidic devices (Krajniak and Lu, 2010, Xian et al., 2013, Atakan et al., 2019, Krajniak et al., 2013, Lee et al., 2014, Gokce et al., 2017). Microfluidic devices allow time-lapse imaging in well-fed animals to measure time-dependent change in fluorescence in neurons for up to 24 hr (Lee et al., 2014). In aging studies, long-term microfluidic devices have been used to grow *C. elegans* and acquire low-resolution images to quantify brood size and monitor locomotion (Xian et al., 2013, Atakan et al., 2019, Rahman et al., 2018). Recently, neuronal branching dynamics and lineage patterns were monitored using a high-resolution imaging platform from an animal growing in a microfluidic chamber over 3 days (Keil et al., 2016). As yet, there is no long-term study tracking sub-cellular events that often need higher resolution and are frequently subject to significant variability. Some examples include organelle distributions and accumulations such as mitochondria and synaptic vesicles. To track a variety of cellular and sub-cellular changes in identified individual animals, we developed simple polydimethylsiloxane (PDMS) microfluidic device to grow *C. elegans* inside micrometer-sized channels and image the same individual neuronal processes throughout its development.

Here, we describe the design and optimization of this microfluidic device and demonstrate its utility in capturing high-resolution fluorescence images of the same posterior lateral touch receptor neurons (TRNs) across 36 hours (hr) of *C. elegans* development. Mitochondrial distribution along the neuronal process changed over time, however, the density of mitochondria was similar across different developmental stages. Our data suggests that this likely occurs through the addition of a new mitochondrion between two mitochondria that are separated by at least 24 μm with the most frequent additions occurring when two mitochondria are 34 µm apart. Our device can also be used to study other events such as the accumulation of synaptic vesicles (SVs) at the synapses, occurring over several days.

## Materials & Methods

### Growth and imaging device fabrication

Our microfluidic device was fabricated using soft lithography of PDMS from SU8 resist features on silicon substrates (Whitesides et al., 2001). The device comprised of two photomasks designed in Clewin software and printed using a high-resolution laser plotter (Fine Line Imaging, USA). The flow layer comprised of a single 10 mm long and 300 μm wide straight channel with one inlet and one outlet reservoirs (Supplementary Fig. 1). The control layer had two independently controlled membranes, through one central wide ‘trapping’ channel and two narrow ‘isolating’ channels. The ‘trapping’ membrane is 2 mm wide to immobilize animals while the restriction membranes are 300 μm wide to keep the worm within the restricted area of the flow channel. The two photomasks were used to produce SU8 masters using UV photolithography for the flow and control layers. The flow layer was fabricated using SU8-2025 (or SU8-2050) spun at 500 rpm for 5 s and 2,000 rpm for 30 s to obtain thicknesses of ∼40 μm (or ∼80 μm). The control layer was created using SU8-2050 spun at 500 rpm for 5 s and 2,000 rpm for 30 s to produce a thickness of ∼80 μm. PDMS (10:1) was prepared by mixing the PDMS base with the curing agent. The PDMS mix was spin coated on the flow layer at 1,000 rpm for 35 s to produce ∼120 μm thickness on the SU8 master and baked at 70 °C for 2 hr. PDMS (10:1) was poured on the SU8 pattern for a 5 mm thick PDMS mold for the control layer and baked at 70 °C for 2 hr. The baked control layer was cut and removed from the SU8 master. Access holes were punched for ‘trapping’ and ‘isolating’ channels and bonded on top of the flow layer using 18 W air plasma for 2 min. The bonded block was finally removed from the silicon substrate, punched through holes for inlet and outlet access, and bonded to a cover glass of thickness 170 μm.

### *C. elegans* strains

We used the *C. elegans* transgenic strains: *jsIs821* [*Pmec-7::GFP::RAB-3*] and *jsIs609* [*Pmec-7::MLS::GFP*] in this study. The strain *jsIs821* expressed GFP::RAB-3, marking synaptic vesicles of the six touch receptor neurons. For time-lapse mitochondria imaging, we used *jsIs609* strain expressing GFP in mitochondria present in six touch receptor neurons.

Strains were maintained on NGM plates using standard protocols (Brenner, 1974). Eggs were transferred to fresh NGM plates and allowed to hatch for two hr at 22 °C. Long-term imaging and mitochondrial transport parameters were calculated at multiple time points up to 36 hr with respect to the hatching time.

### *C. elegans* maintenance inside PDMS device

The flow layer of the device was filled with M9 buffer through the inlet and outlet. The two channels in the control layer were preconditioned with a buffer column under 14 psi pressure of nitrogen gas. When pressurized with compressed nitrogen gas, the column of liquid did not leak to the flow channel due to the presence of the PDMS membrane. A single egg or an early larval stage animal, growing on NGM plates, was pushed into the flow channel within the region between the two ‘isolating’ channels. The two ‘isolating’ membranes were turned on while the central ‘trapping’ membrane was left free. The OP50 bacteria culture was diluted in S medium (1×10^8^ cells/mL), filled with 200 μL micro tips, and connected to the inlet of the flow channel. The outlet was connected with the second pipette with lower food volume. The difference in the height of the food solutions connected to the inlet and outlet channels maintained a constant flow through the flow channel. The smaller overlapping area of the ‘isolating’ channels with the flow channel created a partially closed membrane in the flow channel. The partially closed channel allowed bacteria to flow along the side walls of the flow channel while preventing the animal from crossing the ‘isolating’ membranes. A wide central ‘trapping’ membrane sealed the flow channel completely under similar gas pressure and immobilized animals completely for high-resolution imaging.

### Measuring body length and diameter

Individual animals growing inside the microfluidic devices and on NGM plates were imaged using a brightfield microscope (Olympus IX71) at 4× and 10× magnification. Images were loaded in Fiji (http://fiji.sc/Fiji) and individual worms, with dark features clearly visible on a light background, were traced with segmented lines. The segmented line was drawn from head to tail along the middle of the worm body to measure and quantify body length. The diameter was approximated with a straight line across the body around the vulva location, a region with maximum body width.

### Time-lapse imaging of *C. elegans*

*C. elegans* growing on NGM plates were immobilized using either with 3 mM levamisole on agarose pads or without any anesthetic inside the microfluidic device under the ‘trapping’ membrane for high-resolution fluorescence imaging. Time-lapse fluorescence images were acquired using a 60× (oil immersion and the numerical aperture of 1.4) objective on an inverted microscope (Olympus IX81) equipped with a spinning disc unit (CSU Yokogawa) and a CCD-based camera (iXon, Andor) to visualize the GFP fluorescence of *C. elegans* neurons. Long-term imaging during the development was carried out on the same neuron using multiple images of the neuron at different time points. To maintain animal physiology during long-term imaging experiments, animals were kept free inside the flow channel and in the presence of sufficient bacteria during the intervals between successive imaging sessions.

### Image analysis and quantitation

The physical dimensions of the *C. elegans* body were quantified using brightfield images of animals growing on NGM plates, in 96-well plate cultures, and inside microfluidic devices. Long-term fluorescence images were analyzed in Fiji. For mitochondrial density measurements, multiple image frames covering the complete TRN processes were opened using Fiji. Guided by the features and faint GFP signal present in the neuronal process, the length of the neuronal process was traced between successive mitochondria pair using segmented lines. Every mitochondrial position and the intermitochondrial separations were recorded for individual TRN and specified time points. Numerous total mitochondrial fluorescence envelopes were counted in a given neuronal process from the cell body to the distal tip of the process at each time point. Each fluorescence envelop was segmented to cover all the bright pixels to estimate the average intensity and total area for each mitochondrion. Intermitochondrial distances were quantified as distances in pixels between the centroid of two successive mitochondria in a neuronal process and converted to micrometers. For long-term mitochondrial analysis, the length of the neuronal process of every 3 hr was plotted for individual animal. A linear regression line was fitted to estimate uniform growth rate for each animal. Assuming the uniform growth rate, the length of the neuronal process at every 3 hr time point was normalized, the position of all mitochondria were corrected, and new intermitochondrial distances were calculated for further analysis. Fast time-lapse fluorescence images of the mitochondria were converted to kymographs using multi kymograph plugin in Fiji to quantify the mitochondrial dynamics along the neuronal process. The size of the synaptic accumulation was calculated from the total number of pixels present inside the fluorescence envelope, consisting of all the pixels with fluorescence intensities higher than the background.

### Statistical analysis

Data are presented as the mean ± standard deviation (or standard error of the mean). Histograms are used to show the intermitochondrial distances, gain insight into the shape and variability present in the data, and estimate the probability distribution from the experimental values. We used the Freedman-Diaconis rule to estimate the bin size from the interquartile range (*IQR*) and the number of (*n*) intermitochondrial observations (Freedman and Diaconis, 1981). The bin width was estimated using *n* 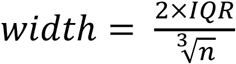, where the *IQR* is defined as the difference between the third and the first quartiles. Statistical tests for significance were carried out using pairwise *t*-tests of unequal variances between the two populations. The significances are presented as a single star (*) and double star (**) wherever p-values are <0.05 and <0.005, respectively.

## Results

### Mitochondrial number along the neuronal process increases with development in *C. elegans* TRNs

Mitochondria are essential organelles involved in energy metabolism and play a vital role in diverse biological processes such as aging and apoptosis (Sheng, 2014, Seervi and Xue, 2015, Sun et al., 2016, Melentijevic et al., 2017). Transport and distribution of mitochondria in neuronal processes and at synapses are critical for the normal physiology of neurons (Morsci et al., 2016, Cai and Tammineni, 2017). Since *C. elegans* are transparent throughout their development, it is feasible to track fluorescently labeled organelles such as mitochondria and SVs in TRNs using high-resolution imaging. *C. elegans* TRNs are a suitable model for high-resolution *in vivo* organelle imaging due to its long and planar neuronal process along the body length present close to the cuticle (Fig. 1*A,B*).

**Figure 1.**
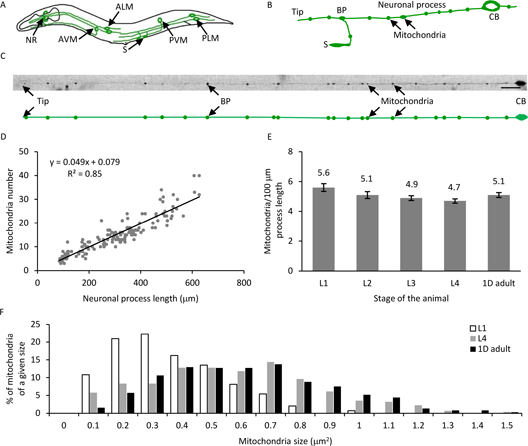
Mitochondria number and process length increase linearly with development. ***A***, Schematic of *C. elegans* with six touch receptor neurons (TRNs). ***B***, Schematic of posterior lateral mechanosensory (PLM) neuron cell body (CB) with a long major neuronal process with a bifurcation of the process at the branch point (BP) to form the synaptic branch that forms synapses (S) along the ventral cord. The mitochondria are indicated with arrows. ***C***, Image of a PLM neuron with 22 mitochondria along the neuronal process. The schematic below the image shows the positions of all 22 mitochondria in the same neuronal process. ***D***, Mitochondria number scales linearly with neuronal process length. Neuronal process length and number of mitochondria of all four larval stages (L1, L2, L3, and L4) and young adult (YA). The trend follows a linear regression with a slope of 4.9 mitochondrial per 100 μm length (total number of animals= 154). ***E***, Mitochondria density remains constant throughout its development. Data represented as mean ± SEM (standard error of mean) for each L1 (n = 30), L2 (n = 25), L3 (n = 36), L4 (n= 33), and YA (n = 30). ***F***, Histogram of the normalized number of mitochondria of a given size represented as a % in the TRNs of L1, L4, and YA stage of animals (n ≥ 148 mitochondria for each stage). The scale bar (c) 20 μm.

To quantify mitochondrial density in *C. elegans* TRNs, we immobilized *jsIs609* animals and imaged the TRNs using mitochondrial matrix targeted GFP expressed in six TRNs for a single time-point analysis during their development. We immobilized the animals on a 2% agarose pad using 3 mM sodium azide and counted mitochondria number, within 10-15 min after the application of the anesthetic, using mitochondrial GFP and length of the neuronal process visible due to some GFP that fills the neuronal process in the posterior lateral mechanosensory (PLM) neurons (Fig. 1*C*). An increase in neuronal process length has been shown to correlate with an increase in mitochondria number across developmental stages (Morsci et al., 2016). We found the neuronal process length increases from ∼100 μm in early larval stage 1 animals (L1) to nearly 500 μm in 1 day adult (1d adult) with a concomitant increase in mitochondria number from approximately 6 to 26 in the posterior TRN (Fig. 1*D*). The bulk of the increase in the length of the neuronal process occurs as the animal grows and occurs after synapse formation at the early L1 stage of the development. The addition of new mitochondria maintains the mitochondrial density of ∼5 mitochondria/100 μm along the neuronal process length throughout development (Fig. 1*E*). This density is similar to earlier reports for this neuron (Morsci et al., 2016, Sure et al., 2018). From the above values, the increase in mitochondria number during development is calculated as ∼6 mitochondria every 24 hr (averaging ∼1 mitochondrion every 4 hr). The area of mitochondria fluorescence in L1, L4, and YA stages of the animal is measured to vary between 0.05-0.95 μm^2^, 0.05-1.66 μm^2^, and 0.05-2.14 μm^2^, respectively (Fig. 1*F*). Young larval animals have a greater number of smaller mitochondria compared to later larval stages or adult animals. Single time point images of *C. elegans* populations of different developmental stages indicate a steady increase in the total number of mitochondria but fail to indicate the location of such addition. To capture individual mitochondrion addition events such that a similar mitochondria density is maintained over development, the same neuronal process needs to be imaged over days under optimal growth and imaging conditions. Since mitochondria are susceptible to cellular stresses (Hill and Van Remmen, 2014, Jovaisaite et al., 2014, Labbadia et al., 2017), it is impossible to image the same animal by repeatedly anesthetizing them for a prolonged period. To reduce the adverse effects of anesthetics and provide a physiological environment, we developed a long-term growth and imaging microfluidic platform.

### Microfluidic device facilitated growth and high-resolution imaging of *C. elegans*

To study the long-term development of individual *C. elegans* while carrying out high-resolution imaging, we developed a microfluidic device to mimic a physiological environment by supplying constant food and immobilizing the animal under a deformable PDMS membrane. To make this microfluidic technology easily accessible, we developed a device with a simple design, few fabrication steps, easy operation, no complex valves, and inexpensive accessories. The PDMS device is bonded on a thin cover glass to facilitate high-resolution imaging on an inverted microscope using lenses of various magnifications up to 100× oil objective.

The device is fabricated as two separate PDMS layers and irreversibly bonded together (Fig. 2*A,B* and Supplementary Fig. 1). The device utilizes PDMS membrane deflections to isolate and immobilize a *C. elegans* hermaphrodite within the flow channel. In this device, *C. elegans* swim freely and grow in a 300 μm wide flow channel in the presence of the bacteria. A pair of partially closed membranes above the symmetric ‘isolating’ channels allows the bacterial suspension to pass through while preventing an individual animal from escaping the flow channel. The individual animal is repeatedly immobilized under a deflected ‘immobilizing’ PDMS membrane. The animal so trapped is imaged for *in vivo* cellular processes beneath the ‘trapping’ channel.

**Figure 2.**
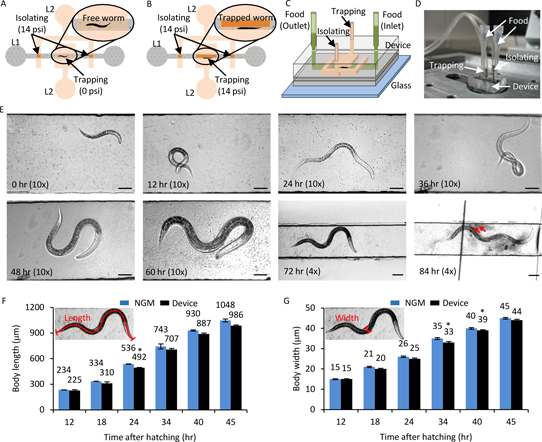
Growth and development of *C. elegans* are unaffected in the long-term growth and imaging device. ***A***, Schematic of the microfluidic device with a flow channel in the bottom layer (L1, grey) and ‘isolating’ and ‘trapping’ channels in the control layer on top (L2, orange). The ‘isolating’ membrane is always kept on (with 14 psi pressure, deflected membrane denoted by dark orange) to restrict the free animal shown in the inset. ***B***, Trap pressure is turned on (14 psi, deflected membrane denoted by dark orange) to immobilize the animal inside the flow channel (inset) during imaging. ***C***, 3D view of the device layout with two pipettes tips (light green) as food inlet and outlet connecting the flow channel (dark green). ***D***, Image of a growth and imaging device connected to compressed nitrogen gas and food supply. ***E***, Images of a *C. elegans* hermaphrodite growing inside the microfluidic device at 0, 12, 24, 36, 48, 60, 72, and 84 hr post-hatching. The animal is fed with OP50 bacteria and isolated inside the flow channel and imaged with 4× or 10× objective. Scale bars are 50 μm, except 100 μm at 72 and 84 hr time points. *jsIs609* animals grown in the device (black) and regular NGM plates (blue) were used for calculating body length (***F***) and body width (***G***). The small inset shows the estimates for body length and body width from worm images. The average values mentioned on top of each bar. Data represented as Mean ± SEM, n = 12 animals. Two-tailed *t*-test, *p*-value<0.01 (*).

Individual animals of the appropriate developmental stage were picked from an NGM plate, using 5 μL M9 buffer, and inserted inside the flow channel in the absence of immobilization or isolation pressure. The animal inside the flow channel was observed using either a 4× or a 10× objective on an inverted microscope. The height of the bacterial solution in the two pipette tips, connected at the inlet and outlet, generated sufficient hydrodynamic flow rates to allow bacterial passage through the flow channel to feed the worm (Fig. 2*C,D*). The deflected membrane in the ‘isolating’ channel restricted animal motion between the two ‘isolating’ channels. Freely moving worms were imaged in brightfield to measure their physical parameters and assess animal health. Animals were completely immobilized in the flow channel under the PDMS membrane using 14 psi pressure in the ‘trapping’ channel. Complete immobilization was required for high-resolution time-lapse imaging of the sub-cellular events in the neuronal processes. Animals were immobilized repeatedly in straight posture along the flow channel sidewalls under the trapping membrane to bring the same posterior lateral TRN under the best focus for imaging the sub-cellular events. Identical animal orientation enabled us to capture mitochondria distribution from the same individual neuron over multiple time points using the inverted confocal microscope equipped with a 60× oil objective (Numerical Aperture 1.4).

### Growth in a microfluidic device does not impede *C. elegans* development

To obtain high-quality time-lapse movies of the sub-cellular events during *C. elegans* development, the health of the animal growing in a microfluidic environment during the entire period should be similar to that of an animal growing on a standard NGM solid media. *C. elegans* growing in a liquid medium with *E. coli* bacteria have been studied previously (Keil et al., 2016, Solis and Petrascheck, 2011). *C. elegans* were found to develop ∼15% slower in liquid culture with adequate bacterial supply but recapitulated all long-term developmental processes such as molting and egg laying (Keil et al., 2016). Since food supply is one critical component for the normal development, we ensured sufficient bacterial supply by adjusting both the concentration of bacteria and the relative bacterial-buffer solution amounts in the two pipette tips attached to the channel entrances. The flow channel containing liquid OP50 showed no contamination over 84 hr (4 days) of *C. elegans* growth (Fig. 2*E*). The pipette tips are easily replaced with new sterile tips freshly filled with OP50 bacteria.

To characterize *C. elegans* development in our microfluidic device, we measured the body length and body diameter of the animals grown in our microfluidic chamber and compared the values with body sizes from animals grown on NGM solid media. *C. elegans* have been shown to grow slightly slower in liquid culture (Keil et al., 2016). Our data show that animals growing in the microfluidic device are on average shorter and thinner compared to those grown on the NGM media (Fig. 2*F,G*). The hermaphrodite growing inside the device laid eggs after 84 hr, which hatch into larvae.

### Microfluidic device facilitates repeated immobilization and time-lapse imaging of same neuronal process in *C. elegans*

Mitochondria in neuronal process display distinct transport characteristics; constant velocity away from (anterograde) and towards (retrograde) the cell body, intermittent pause, and bi-directional movement (O’Toole et al., 2008, Cai and Sheng, 2009, Schwarz, 2013, Welte, 2004, Hancock, 2014, Sure et al., 2018). To quantify *in vivo* mitochondrial transport characteristics from the *C. elegans* TRN processes, we immobilized animals in the flow channel using 14 psi on the trapping PDMS membrane (Fig. 3*A*). A slow progressive immobilization allows animals to be laterally oriented, facilitating high-resolution imaging in the PLM neurons.

**Figure 3.**
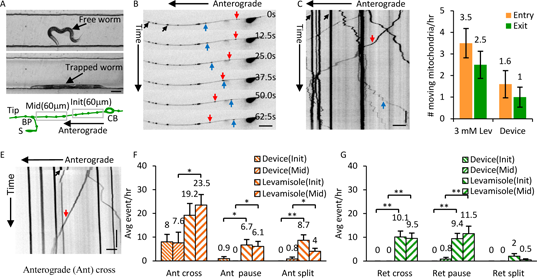
High-resolution transport imaging of mito::GFP in device immobilized *C. elegans* neurons. ***A***, Image of a free and an immobilized larval 4 (L4) stage *C. elegans* in the flow channel. The schematic shows the initial 60 μm from the cell body and middle of the neuronal process (120 μm from the branch point towards the cell body) that is imaged at high-resolution. ***B***, Montage of six frames of a PLM neuron with mito::GFP fluorescence from time-lapse imaging acquired using a 60× oil objective (1.4 NA). ***C***, Kymograph of 400 frames acquired at 2 frames per second (fps) for 200 seconds. Anterogradely moving (red arrow), retrogradely moving (blue arrow), and stationary (black arrow) mitochondria are indicated in the images. ***D***, Number of mitochondria entering the neuronal process (entry) and leaving the process (exit) per hr measured in animals immobilized with 3 mM levamisole (n = 35 animals) and inside the microfluidic device (n = 22 animals). None of the pairs are statistically different (*p*-value ≥ 0.055). ***E***, Kymograph representing an anterograde moving mitochondrion (red arrow) crossing a site of the bleached mitochondria (black arrow). ***F***, The average anterograde events per hr (cross, pause, and split) are measured at the initial (Init, 60 μm from the cell body) and the middle (Mid, a region 120 μm from the branch point towards the cell body) of the neuronal process (schematic is shown in panel ***A***) for both device (Device, n > 8 animals) and anesthetic (Levamisole, n > 29 animals) immobilized L4 animals, respectively. ***G***, The average retrograde events are measured by using the device and anaesthetic immobilized animals. Data represented as Mean ± SEM. Statistics are calculated using a two-tailed *t*-test, *p*-value < 0.05 (*) and < 0.005 (**). The scale bars are 100 μm (***A***), 10 μm (***B*** and ***E***), and 5 μm (***C***). The vertical bars are 20 s (***C*** and ***E***).

To assess the effect of photobleaching and changes in auto-fluorescence due to repeated immobilization, we repeatedly immobilized *C. elegans* at two different time intervals; either every 1 hr (trapping every 1 hr) or every 6 hr (trapping every 6 hr), inside a microfluidic device for up to 12 hr under “immobilizing” membrane using 14 psi pressure. For both conditions, the same PLM neuron (either L or R) of the animal was only imaged at 0, 6, and 12 hr time points. The ratio of the mitochondrial fluorescence to the background fluorescence (M/B ratio) was calculated at different time points. The drop in the fluorescence ratio for the two immobilization cycles was statistically insignificant (2.07 ± 0.106, 1.80 ± 0.192, *p* = 0.06) (Supplementary Fig. 2). Using repeated imaging of *C. elegans* in our microfluidic device, we were able to obtain good fluorescence signals from mitochondria compared to the background autofluorescence.

We further characterized the effect of immobilization of an individual *C. elegans* on the variability in the position of each stationary mitochondrion that arises just from the immobilization process and small changes in the posture of the animal. Positions of mitochondria were used to calculate the intermitochondrial distances and estimate the local compression and expansion of the neuronal process between multiple adjacent mitochondrial pairs. We, therefore, imaged the same TRNs every 5 min for 15 min by repeatedly trapping the animal under the ‘immobilizing’ PDMS membrane and calculated intermitochondrial intervals for each immobilization. Intermitochondrial distance values (L) between identical mitochondria pairs from the same TRN processes imaged at different time points were compared (ΔL) to estimate the extent of expansion or compression between each mitochondria pair (ΔL/L). Only stationary bright mitochondria were considered for this calculation. Most mitochondrial positions remain largely unchanged and intermitochondrial distances were used to measure the compression or expansion percentages between each pair of mitochondria (Supplementary Fig. 3*A*). The compression and expansion percentages were distributed symmetrically and represented as positive (compression) and negative (expansion) values, respectively. The ΔL/L values were distributed around +1.79 % and -2.01 % (2.4 ± 0.17 μm and 2.4 ± 0.19 μm, mean ± SEM) (Supplementary Fig. 3*B*). Thus, the errors arising from compression and expansion due to immobilization does not significantly alter the measurement of intermitochondrial distances.

### Device-immobilized animals show lower mitochondria flux and turnover compared to anesthetic immobilized animals

We examined mitochondria number and their dynamics in NGM grown and device grown animals. The density of mitochondria in the larval stages and the YA animals grown in NGM and microfluidic devices are similar. Additionally, the inter-mitochondrial intervals in NGM grown animals and device grown animals are very similar. To examine the movement and turn over of mitochondria from both device and anesthetic immobilized animals, we carried out high-resolution time-lapse imaging of TRNs. The acquired movies were converted into kymographs to analyze mitochondrial transport and positions (Fig. 3*B,C*). Device immobilized animals regained normal body movements within 10 min of release from pressure. Mitochondria are classified as (1) stationary if their displacement is less than 3 pixels in 5 consecutive frames and (2) moving if their displacement is equal to or more than 3 pixels in less than 5 consecutive frames. Mitochondria move anterogradely away from the cell body or retrogradely towards the cell body. The movement close to the cell body was captured for 20 min at 3 fps to quantify all anterograde and retrograde events in both anesthetized animals and device immobilized L4 stage animals, both grown on NGM plates until imaged. The average number of mitochondria entering the neuronal process from the cell body was about twice the total number of mitochondrial exit events into the cell body (Fig. 3*D*). The average number of entry and exit events were lower but not statistically different (*p*-value > 0.5) in the device immobilized animals compared to the anesthetized animals. In device immobilized animals, the rate of mitochondrial entry into the process is twice that of mitochondrial exit as well (Fig. 3*D*).

Using fast time-lapse imaging of the individual TRNs, at a speed greater than 4 fps and for a total time ≥ 150 s, we analyzed turnover rates of moving mitochondria across photobleached stationary mitochondria. Upon photobleaching stationary mitochondria, the motion of a small and faint moving mitochondrion can easily be tracked across the site of the stationary mitochondria. In a simple cross event, the small mitochondria moved across the photo-bleached site without contributing fluorescence to the stationary mitochondria (Fig. 3*E*). While moving mitochondria are identified as contributing to photobleached stationary mitochondria by either pausing and/or splitting a portion of their fluorescence leading to a recovery in the large stationary mitochondria fluorescence (Supplementary Fig. 4*A-C* and Supplementary Table 1). Sometimes paused mitochondria filled the entire region of the stationary mitochondria before photobleaching and sometimes they merely paused and continue to move. Both anterograde and retrograde moving mitochondria were found to contribute to the turnover of stationary mitochondria. On an average, more events were observed in anesthetized animals compared to the device-immobilized animals (Fig. 3*F,G*). Both the device and levamisole immobilized animals show a higher number of anterograde movements across stationary mitochondria as compared to that measured in the retrograde movements. We did not observe a stationary mitochondrion to split into smaller mitochondria in any of our movies during the entire 4.6 hr of imaging either under anesthetic or immobilized in a device. The fluorescence envelops of the moving mitochondria were smaller than the stationary mitochondria. We sometimes observed tubular-shaped moving mitochondria in the first 60 μm of the neuronal process. The high amount of stress introduced to the animal in an anesthetic assay perhaps accounts for the increase in the mitochondrial activity which may cause a high turnover of mitochondria along the neuron.

### Mitochondria are uniformly added to the *C. elegans* neuronal processes growing in microfluidic devices

Mitochondria are added in a growing neuron after synapse formation to maintain a constant mitochondrial density. For example, the number of mitochondria and total process length respectively increased from 7.8 ± 0.41 and 125 ± 6.07 μm in the L2 stage to 20.1 ± 1.44 and 335 ± 15.89 μm in L4 animals when measured using mitochondrial matrix targeted GFP inside the microfluidic device (Supplementary Fig. 5). The neuronal process grows at the rate ∼10 μm/hr and the mitochondria number increases ∼0.6 mitochondria/hr, numbers very similar to those seen in animals grown on NGM plates (see above). Although the population statistics indicate average growth rates, it does not indicate the site where new mitochondria were added.

To identify the location where the new mitochondria are added as the animal is growing, we imaged the same PLM neuron every 3 hr for approximately 36 hr by repeatedly trapping the animal in our microfluidic device. The images were used to reconstruct the neuron and the location of each mitochondrion was plotted for every imaged time point. The total mitochondria number increased linearly with the imaging time. Both, total mitochondria number and neuronal process length at 24 hr were significantly higher than the initial time point (Fig. 4*A*). We chose the cell body, branch point, and the neuronal process tip as the fiduciary markers to examine if there was any bias in the growth of the neuronal process and during the new mitochondria addition events along the neuronal process (Fig. 4*E*). The ratio of the neuronal process lengths between the cell body to the branch point and the branch point to the neuronal process tip remains the same as the neuron and the animal grows suggesting proportional growth along the neuronal process (Fig. 4*B*). The uniform increase in neuronal process elongation (Fig. 4*C*) is consistent with the earlier studies that show uniform membrane addition in the growing neuron process (Popov et al., 1993, Chen et al., 2014). Over 36 hr of imaging, the average total neuronal process growth rate was calculated to be ∼9 μm/hr (n = 8 animals). We calculated the ratio of the mitochondria number present on the neuronal processes between the cell body and the branch point to the mitochondria number between the branch point and the neuronal tip. The ratio of the mitochondria number remains the same over 24 hr (Fig. 4*B*), which suggests that new mitochondria are likely added uniformly along the process length.

**Figure 4.**
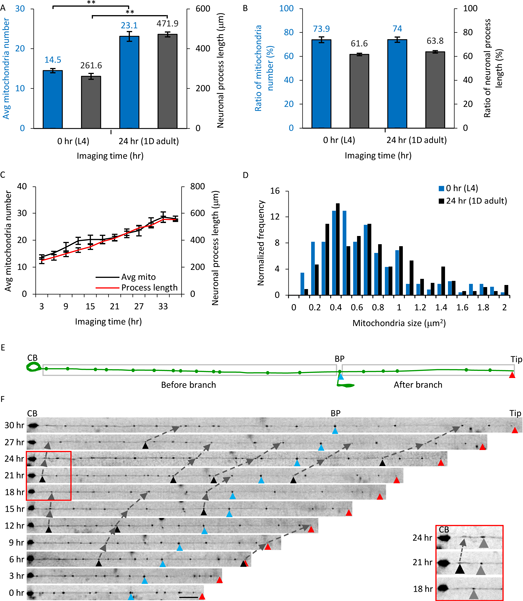
Total number of mitochondria increases linearly in TRNs of the animals growing inside microfluidic devices. ***A***, The average number of mitochondria and the neuronal process length increases starting 0 hr (L4) to 24 hr (YA) inside the microfluidic device. ***B***, The ratio of the mitochondria number and the process length was unchanged over this time period. The ratio is calculated for the values from the cell body to the branch point and from the branch point to the neuronal tip. ***C***, Time course of the mitochondrial number and neuronal process length from 8 animals imaged every 3 hr in microfluidic devices beginning at L4. Data represented as mean ± SEM. Statistics are calculated using a two-tailed *t*-test, *p*-value<0.005 represented as ‘**’ (number of animals=8). ***D***, The relative frequency distribution of all the mitochondria size at 0 and 24 hr (n=8 animals). A two-tailed *t*-test was calculated between the two-time points for statistics (*p*-value = 0.92). ***E***, A schematic of the TRN showing the neuronal process connecting the cell body (CB) and the tip (red triangle) with the branch point (BP, blue triangle) as fiduciary markers. The branch point is used to estimate the total mitochondria number and neuronal process length between the cell body and the branch point and between the branch point and the end of the neuronal process. ***F***, The same animal was trapped inside the microfluidic device every 3 hr and imaged at high-resolution to image mitochondria in the same TRN. Fluorescence images were straightened and raw images are displayed. The fiduciary markers of the cell body, branch point, and neuronal end are used as references. Addition of new mitochondria are labeled with black triangles and followed in two successive frames using dashed lines. Scale bar 20 μm. The red box and the inset show a section of the neuronal process at 18, 21, and 24 hr time points. A new mitochondrion (black triangle) is added in between the cell body and the stationary mitochondria (gray triangle). The distance between the cell body and the stationary mitochondria is increasing with time due to the growth of the neuronal process.

To capture every mitochondrion, we reconstructed the whole neuronal process from a series of fluorescence images at multiple time points and analyzed various parameters such as mitochondrial number, size, and the intermitochondrial interval between each adjacent pair. Approximately 20% of mitochondria move in any given frame in the TRNs with an average velocity of approximately 200 nm/s (Fatouros et al., 2012). The moving mitochondria are less than ∼1 μm in diameter. Stationary mitochondria are bigger in size but drift gradually over time as the animal grows and the neuronal process commensurately elongates proportionately. Small moving mitochondria have low fluorescence intensity and appear randomly along the TRN processes in our long-term images due to large time intervals between successive frames. The size of the stationary mitochondria in the neuronal process remains unchanged over longer time scales e.g. 24 hr (Fig. 4*D*). We consider a pair of mitochondria to be identical and stationary if the position of each mitochondrion from the cell body, branch point, and tip of the neuron changes proportionately to the average growth rate of the neuron for that animal (typically ∼27 μm per 3 hr). The neuronal processes of each individual TRN at multiple time-points are aligned with each other from the cell body where the cell body, the branch point, and the neuronal process tip act as reference points (Fig. 4*F*). With uniform growth of the neuronal process, the fiduciary markers (BP and tip) and all the mitochondria move further away from the cell body with time (Supplementary Fig. 6*A*). For every mitochondrion, the ratio of the new position to the old position was 1.1 (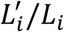 where, 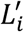 and *L*_*i*_ are the new and old position of the *i*^*th*^ mitochondria with respect to the cell body), correlating with the 10% increase in the total neuronal process length over 3 hr. The uniform increase and the gradual shift in the stationary mitochondria positions are independent of the alignment of the neuronal process with respect to the cell body or the tip of the neuron (Supplementary Fig. 6*B*).

To identify an event of a new mitochondrion addition, we find a mitochondrion between a pair of pre-existing stationary mitochondria with higher fluorescence than the background intensity from the neuronal process. The new mitochondrion must stay in the same location for at least two successive frames (i.e. greater than 6 hr). The addition of a new mitochondrion was identified using manual inspection of the relative mitochondrial position and geometrical parameters after aligning the axons with respect to the fiduciary markers (Supplementary Fig. 6).

The number of mitochondria addition events was calculated for every equal third of the neuronal process i.e. initial, middle, and end. The number of events in each third of the neuron was similar 33.3 ± 3.24% (n = 29), 37.5 ± 2.61% (n = 32), and 29.1 ± 3.31% (n = 24) with *p* ≥ 0.07 (Supplementary Fig. 7). We see that each new mitochondrion added has an approximate size of 0.47 ± 0.07 μm^2^ (mean ± SEM) when first identified and grew to 0.68 ± 0.11 μm^2^ (n = 4 animals and n = 24 new mitochondria addition events) after 3 hr. The gain in mitochondrial size can come from contributions from either the moving mitochondria or in situ biogenesis. Moving mitochondria along the neuronal processes are likely to initiate a new location and subsequently gain mitochondrial material by fusion-fission of moving mitochondria to form a larger stationary mitochondrion over time in the neuronal process. Using a long-term imaging platform, we were able to identify the addition of new mitochondria in an identified neuronal process.

### Increased intermitochondrial distance leads to the addition of a new mitochondrion in *C. elegans* TRNs

The uniform growth in the neuronal process increases the intermitochondrial distance between each adjacent pair of mitochondria. This growth of the neuron occurs along with new mitochondria addition to maintain density. To characterize changes in the mitochondria density and identify the site of the new addition, we calculated the intermitochondrial distances between each pair of mitochondria (center-to-center distance along the neuronal process length) in the all eight L4 stage animals (Supplementary Fig. 8). Several cellular possibilities can contribute towards the mitochondria addition events and the intermitochondrial distance as measured from the new mitochondrial pairs (Supplementary Fig. 9). (1) Mitochondria are added near the cell body. (2) Mitochondria are added at the end of the neuron. (3) A new mitochondrion is added when the intermitochondrial distance between adjacent mitochondria increases beyond a threshold. In either possibility (1) or (2) the intermitochondrial distances will show two peaks, one at short and another at long intermitochondrial distance intervals corresponding to additions in one region of the neuron and with the increasing distance between other mitochondria due to axon growth. In the third situation, as the mitochondria move apart due to increasing neuronal process length, the entire intermitochondrial distance distribution will shift to higher values (resulting in a low number of events for the short intermitochondrial distance values). According to the third situation, a new mitochondrion will be added in between the old mitochondria pair and reducing the possibility to have long intermitochondrial distance values. Hence, we expect to observe a small reduction in shorter-intermitochondrial distances but a relatively unchanged longer-intermitochondrial distance. We hypothesize that the addition occurs when the intermitochondrial distance between adjacent mitochondria increases beyond a threshold (orange, Fig. 5). A separate pair of mitochondria moving up to 25 μm, still below the threshold, does not dock a new mitochondrion over the same timescale (gray, Fig. 5*A,B*).

**Figure 5.**
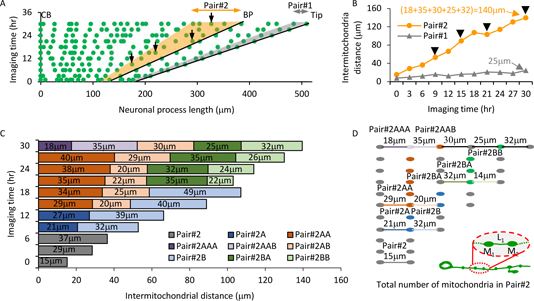
Addition of new mitochondria during the long-time imaging of the animal growing in the microfluidic device. ***A***, Map of mitochondrial position from an individual *C. elegans* PLM neuron after 36 hr of hatching. Solid lines represent all three locations CB, BP, and Tip of the neuronal process. The shaded area in gray (Pair#1) is a region in the neuronal process where no mitochondrion is added in 30 hr of imaging between adjacent pairs. The shaded area in orange (Pair#2) represents regions of the same neuronal process where four new mitochondria are identified between the adjacent pairs at different imaging times. The black arrows represent locations of all the four new mitochondria. ***B***, Intermitochondrial intervals at different imaging times of two different pairs of stationary mitochondria (Pair#1 and 2) from an individual TRN process. The arrows indicate the time-points when a new mitochondrion was added between the two adjacent mitochondria along the TRN process. The values indicate the total distance between the parent pairs at 30^th^ hr of imaging, upon adding all the intermediate intermitochondrial intervals formed due to new mitochondria additions. ***C***, The bar represents the mitochondria lineage for Pair#2 measured from time-lapse imaging. Upon the addition of the new mitochondrion, each pair of mitochondria is labeled as new and represented by two new colors (light and dark). The numbers on each bar indicate the intermitochondrial distances for every mitochondria pair. The short intermitochondrial distances in few time points arise due to local compression/expansion of the neuronal process and are considered to be artifacts of imaging due to animal posture. ***D***, Schematic of the mitochondria lineage in Pair#2. The gray dots on the right represents the original mitochondria pair (not to the scale) while the new color dot indicates the location of the newly added mitochondria. Whenever a new mitochondrion is added, the intermitochondrial distances between new pairs are indicated by a line with a color similar to the color of the bars. The inset shows the schematic of a growing PLM neuron during the development of *C. elegans*. The cell body is on left and the neuronal process tip is on right. The inset also shows two mitochondria (M_1_ and M_2_) separated by length L_1_.

To study the influence of mitochondria addition on a growing neuronal process, we compared the intermitochondrial distances obtained from the images at 0 hr and 24 hr of eight different animals at an early L4 and the young adult stages, respectively (Fig. 6*A*). After 24 hr, the population average of the intermitochondrial distances between adjacent mitochondria increased from 17 ± 0.7 μm to 19 ± 0.6 μm (mean ± SEM, n = 8 animals). The addition of new mitochondria over 24 hr caused a more skewed distribution as compared to 0 hr time point for the intermitochondrial distance (skewness and kurtosis measures increased from 0.814 and 0.341 at 0 hr to 0.894 and 0.798 at 24 hr, respectively). The larger values for the skewness and kurtosis measures at 24 hr indicate a strong heavy- and right-tailed distribution, indicative of non-random mitochondria position as the neuronal process grows. Over the same 24 hr period, the average values for the first two bins (% of adjacent mitochondria separated by 3 μm and 6 μm) reduced from 7 ± 1.03 % and 9 ± 1.27 % to 3 ± 1.17 % and 7 ± 1.01 % (*p*-values = 0.02 and 0.17), respectively. No other bin in the normalized plot has a statistically significant difference in intermitochondrial distances at the end of this imaging period. Although, elongation of the neuronal process causes mitochondria to move apart (as indicated by the significant reduction in the smaller inter-mitochondrial intervals), the addition of new mitochondria ensures an insignificant increase in the larger intermitochondrial values. Although mitochondria undergo dynamic membrane fission/fusion and active transport (Sheng, 2014), their addition and positioning along the TRNs were surprisingly found to be regulated as reflected by the highly skewed distribution of the intermitochondrial distance values. In contrary to our observation, a neuron with randomly localized mitochondria, the intermitochondrial distances from each neighboring pair will follow an exponential distribution.

**Figure 6.**
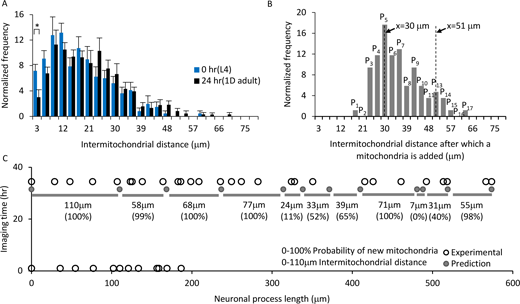
Experimental probability distribution can predict addition of new mitochondria in a growing neuronal process length. ***A***, Normalized number of intermitochondrial distances at 0 hr (L4) and 24 hr (YA) after the start of imaging. Two-tailed t-test, p-value<0.05 (*). ***B***, Normalized number of mitochondria addition is calculated from experimental long-term images of individual TRNs (n=8 animals started at the L4 stage, 85 new mitochondria additions). The probability of new mitochondria added for intermitochondrial distance x = L is shown as P_L_ and can be estimated by summing (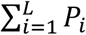 where *i* ≤ *L*). ***C***, The predicted (dark circle) and experimental (open circle) location of mitochondria at 0 and 33 hr images of an individual animal. All the 12 positions at t = 33 hr are predicted using the locations measured at t = 0 hr and total neuronal process elongation values. The intermitochondrial distances (gray bars with the values) are used to predict the addition of new mitochondria between each pair at 33 hr. The probability values for finding new mitochondria within the predicted pairs are represented in percentages.

The intermitochondrial separation initiates the formation of new mitochondria forming two independent pairs of mitochondria, one of the pair (Pair#1A) is formed with the old mitochondria close to the cell body and second pair (Pair#1B) is formed with the old mitochondria far from the cell body (Supplementary Fig. 10*A,B*). The number of newly added mitochondria from all eight animals were normalized to the total number of n = 85 mitochondria addition events, which shows a distribution with an average of 34 ± 1.0 μm (mean ± SEM) (Fig. 6*B*). We identified all new mitochondria pairs to estimate the average distance for Pair#1A and Pair#1B as 17.1 ± 0.8 μm and 21 ± 1.2 μm (mean ± SEM), respectively for all n = 85 events (Supplementary Fig. 10*C,D*). The new mitochondria addition events are uniformly distributed along the neuron (Supplementary Fig. 10*E*). Consistent with the third possibility we observed 85 events where two adjacent mitochondria had an inter-mitochondrial interval >24 μm where a new mitochondrion position appeared and remained for 6 hr or more. The addition of new mitochondria occurs uniformly throughout the neuron and maintained a similar average intermitochondrial distance over a day of long-term imaging.

### Mitochondria positions can be estimated from the experimental probability distribution function

Using all the individual events where a new mitochondrion was added, we calculated the experimental probability distribution of new addition events and predicted possible locations for the addition of new mitochondria using the probability distribution. The normalized plot of intermitochondrial distance (Fig. 6*B*) provides us the experimental probability distribution with a sum of all the values (∑ *P*) = 100%. For any pair of mitochondria, we calculate the intermitochondrial distance (*x* = *L*) and add normalized numbers of mitochondria addition for all the intermitochondrial distances lower than the current value to assign the probability (*P*_*L*_) of the addition of new mitochondria between a given adjacent pair (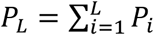, where *i* ≤ *L*). In order to test our probability distribution values, we imaged a neuronal process length (L_0_ = 186.7 μm) with total mitochondria (N_0_ =12) at time t = 0 hr and predict that the hypothetical neuronal process grows to a length (L_33_ = 573.7 μm) at time t = 33 hr after the start of imaging. This gave us 12 images that were collected every 3 hr (Fig. 6*C*). For the ‘predicted’ neuron, we calculated the neuronal process elongated steadily with an experimental rate of 11 μm/hr, similar to the average neuronal process elongation rate of ∼9 μm/hr. Using the above parameters, we estimated the neuronal process (with an average intramitochondrial distance of 17 ± 2.8 μm at t = 0 hr) to grow ∼11 μm/hr for 33 hr leading to an average intermitochondrial interval of 52 ± 8.7 μm, without the addition of new mitochondria (Fig. 6*C*). When the separation between adjacent mitochondrial pairs increased beyond 24 μm, we predicted the addition of new mitochondria for an intermitochondrial separation value by adding all the experimental probability values (Fig. 6*B*). For a large value of the intermitochondrial distance, we added all the probability values (all the bars add up to 100%). For higher values of the probability, we predicted the addition of new mitochondria which matched with the experimental observations (Fig. 6*C*).

This technology allows us, for the first time, to track individual neurons to assess the dynamics that occur over slow timescales of intracellular organelle positioning across development. Using our long-term imaging device, we were able to extract the experimental probability distribution for organelle distribution and use the values to predict the location of a new mitochondrion addition between a pair of adjacent pre-existing mitochondria in a growing neuron. Multiple factors could determine the addition of new mitochondria such as the ratio of molecular motors (Sure et al., 2018) (Souvik M *et al.* submitted with this article), local cytoskeleton (Sood et al., 2018), local calcium concentration (Wang and Schwarz, 2009b), new docking sites (Cai and Sheng, 2009), and/or reactive oxygen species (Debattisti et al., 2017, Liao et al., 2017).

### Our long-term device can be used to study synaptic development in an individual animal

We wanted to test whether our device could be used to image other developmental processes to examine its general utility for other types of biological processes. To perform high-resolution time-lapse imaging of sub-cellular organelle accumulation events in *C. elegans* across developmental stages, we captured fluorescence images of GFP::RAB-3 accumulation at the ventral PLM synapses from the same individual *jsIs821* animals (Kumar et al., 2010, Nonet, 1999, Mahoney et al., 2006). We immobilized each animal using our imaging platform under 14 psi pressure inside the flow channel. Slow progressive immobilization provides a suitable lateral orientation of *C. elegans* and the best view of the PLM synapses (Supplementary Fig. 11). We immobilized the same animal in lateral orientation to image both PLM synapses at L2 (16 hr), L3 (22 hr), L4 (38 hr), and 1d adult (68 hr) from the time the eggs were hatched. High-resolution time-lapse images were analyzed for synapse sizes to estimate synapse growth at different developmental stages. The amount of GFP::RAB-3 accumulation shows a gradual increase similar to the prior average signal measured from multiple *C. elegans* animals at different developmental stages (Mondal et al., 2011). This increase is thought to arise from the gradual increase in the net anterograde flux of synaptic vesicles (Mondal et al., 2011). Our imaging platform captured a robust increase in the accumulation of GFP::RAB-3 at the synapses of the same animal over multiple developmental stages.

## Discussion

*C. elegans* TRNs grow from approximately 100 μm at the L1 stage to 500 μm in 1d adult stage that results in the addition of approximately 20 new mitochondria to maintain a constant mitochondrial density of ∼5 mitochondria/100 μm of the neuronal process length. The density measured in adult animals is found to be very similar to that previously reported in YA animals (Morsci et al., 2016, Sure et al., 2018). Conventional anesthetic assays can have detrimental effects on animal physiology and cannot easily be used for long-term imaging of the same animal to follow slow processes such as mitochondria addition during neuronal process growth. In this work, we developed a microfluidic device that can be used to grow *C. elegans* throughout its development without significant adverse effects on its growth and phenotype. Its easy operation enables continuous imaging of the same animal inside the microfluidic environment to quantify *in vivo* sub-cellular process of the neuron throughout development in the narrow device geometry. Mitochondria in the TRNs were found to show lower flux in the microfluidic device compared to anesthetic immobilization. These observations are similar to the lower flux of synaptic vesicles when immobilized in microfluidic devices compared to anaesthetic immobilization (Mondal et al., 2011). Using long-term imaging over a period of 36 hr, we identified mitochondria addition and changes in intermitochondrial distances in *C. elegans* TRNs during its elongation after synapse formation. Using our microfluidic device platform, one can monitor slow processes, here the addition of mitochondria. Slow entry and exit, and limited turn-over of mitochondria identified in our imaging platform suggests that the neuron retains positions of old mitochondria and adds mitochondria at new locations along the growing process between pre-existing mitochondria during development. A new location for mitochondrion addition appears to emerge whenever the average separation between two nearest neighbors is at least 24 μm.

We have identified the experimental probability distribution of the addition of a new mitochondrion in a growing neuronal process *in vivo* over 24 hr. The distribution for the intermitochondrial distances becomes more skewed and the mean increases with the development of the animal (Fig. 6*A*). With age, the number of closely spaced mitochondria is reduced, decreasing the fraction of mitochondria with lower separation between them. During neuronal process elongation and addition of a new mitochondrion, the newly added mitochondria are small in size and have lower integrated fluorescence intensity that grows over time (Supplementary Fig. 6). There are two possible mechanisms for the addition of new mitochondrion that remains a subject for further studies; 1) a small mitochondrion enters from the cell body, transports along the neuronal process, identifies a large intermitochondrial separation, and anchors within the “mitochondria-in-demand” region and 2) a local mitochondrion fission event causes a new small mitochondrion to eject from a nearby large mitochondrion, transports along with a shorter distance, and anchors within the “mitochondria-in-demand” region. However, we have not observed a large stationary mitochondrion show such fission in our imaging. All events appear to be initiated by moving mitochondria undergoing fusion or fission with a stationary mitochondrion. Thus, the addition of new mitochondria likely arises from the stalling of moving mitochondria at a defined location. The stalled mitochondria appear to enlarge in size potentially through fusion with material from passing moving mitochondria. We believe that time-lapse imaging of mitochondria pairs greater than 34 μm apart, identified from our long-term time-lapse imaging platform, could shed light on mechanisms of mitochondrial addition. Time-lapse imaging of fission mutants and animals with transport defects can help delineate the underlying process/molecular players necessary for the new mitochondria addition events.

Our device can be used at different time intervals. For studies that require more frequent imaging of the same animal, we immobilized the same animal every hour for 12 hr (Supplementary Fig. 2). With 1 hr frequent imaging interval, we observed greater photobleaching and a more slowly growing animal beyond 12 hr. Intermittent long-term imaging separated by ∼6 hr for 52 hr shows a steady increase in synaptic accumulation of GFP::RAB-3 when measured from an individual animal growing inside the microfluidic device. The amount of accumulation was slightly higher in the device grown animals as compared to animals grown on NGM plates. This increase compared to animals grown on NGM plates may arise from additional mechanical stimulation provided by the deflected membrane pressing against the animal. Despite these caveats, our microfluidic platform can be utilized for various studies to assess long-term cellular or sub-cellular dynamics in *C. elegans*. Similar technologies can be used for other model organisms by modifying the channel geometries.

## Acknowledgments

The *C. elegans* strain jsIs821 were made in Mike Nonet’s Laboratory. We acknowledge Anyaa Mittal and Arpan Agnihotri for their help with microfluidic device fabrication and growing *C. elegans* inside microfluidic devices. We are grateful to Gautam Menon for useful discussion on mitochondria transport imaging and image analysis. We would like to thank Sangeetha Iyer (Perlara Labs), Amruta Vasudevan, and Souvik Modi for suggestions on our manuscript. We thank the CIFF imaging facility, NCBS for use of the confocal microscopes supported by the DST – Centre for Nanotechnology (No. SR/55/NM-36-2005). We thank research funding from DBT (SPK), DST (SM), DBT (SM), CSIR-UGC (JD), spinning disc supported by DAE-PRISM 12-R&D-IMS-5.02.0202 (SPK and Gautam Menon), and HHMI-IECS grant number 55007425 (SPK).

