## Supplementary information for "Tracking mitochondrial density and positioning along a growing neuronal process in individual *C. elegans* neuron using a long-term growth and imaging microfluidic device"

**Supplementary Figure 1:** Schematic of the flow and control layers.

**Supplementary Figure 2:** Effect of photobleaching in long-term mitochondrial imaging.

**Supplementary Figure 3:** Effect of repeated immobilization on intermitochondrial distances between stationary mitochondria present on the TRNs.

**Supplementary Figure 4:** Kymographs of L4 stage *C. elegans* showing anterograde moving mitochondria across bleached mitochondria.

**Supplementary Figure 5:** Mitochondrial density in L2 and L4 animals measured from microfluidic device immobilization.

**Supplementary Figure 6:** Time-lapse images of the neuron from an individual animal.

**Supplementary Figure 7:** Normalized number of mitochondria addition events at the initial, middle, and tip of the neuronal process.

**Supplementary Figure 8:** Intermitochondrial distance measured from L4 stage animals immobilized in the microfluidic chip.

**Supplementary Figure 9:** Schematic representation for mitochondria addition events between pre-existing mitochondria in a growing neuronal process and corresponding changes in inter-mitochondrial intervals.

**Supplementary Figure 10:** Intermitochondrial distance statistics when new mitochondria are added in between a pair of old mitochondria.

**Supplementary Figure 11:** High-resolution imaging of GFP::RAB-3 in a developing *C. elegans* inside the microfluidic device.

**Supplementary Table 1:** Turnover of mitochondria events at stationary mitochondria in the PLM neurons of L4 animals immobilized with 3 mM levamisole.

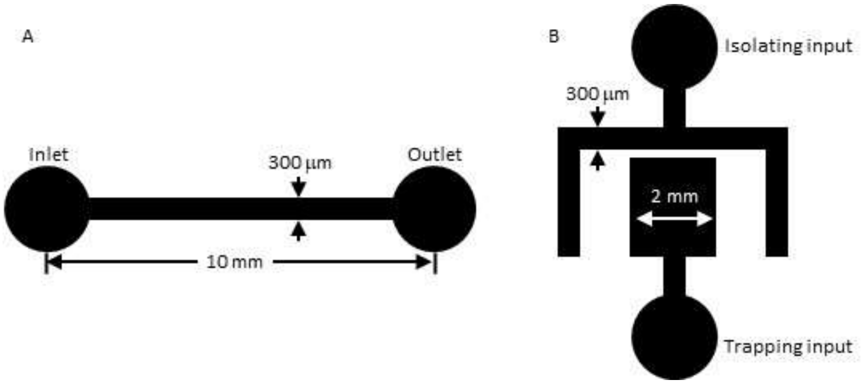


**Supplementary Figure 1:** Schematic of the flow and control layers. ***A***, The schematic for the flow layer showing the dimensions of the channels. ***B***, The schematic of the control layer with the channel dimensions. The circular pads for the punches and fluidic connections are 2 mm in diameter.

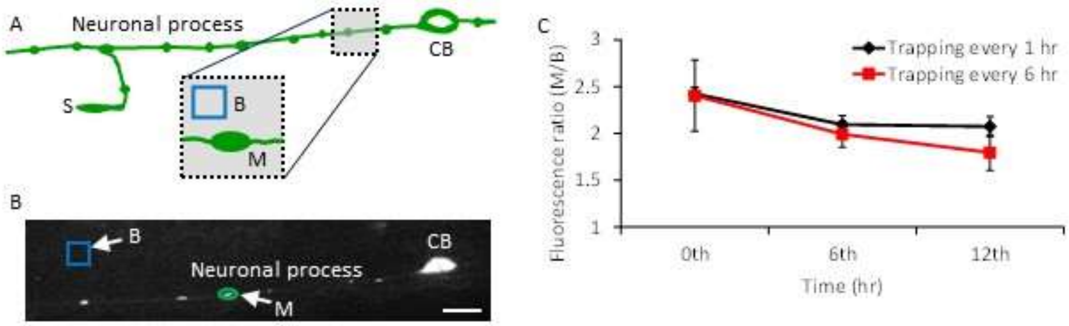


**Supplementary Figure 2:** Effect of photobleaching in long-term mitochondrial imaging. ***A***, Schematic of the TRN neuron. The inset shows a single mitochondrion and a 5×5 μm box to calculate the average intensity values for a mitochondrion (M) and the background (B). ***B***, Image of a single neuron at zero time showing a single mitochondrion (M) and the background box (B) on the worm body. The scale bar is 10 μm. ***C***, Fluorescence ratio of the mitochondria to background intensity (M/B) is calculated from the images of the same animal captured at 0th, 6th, and 12th hr time points. The animals were immobilized at two different time intervals of every 1 hr and every 6 hr, respectively. Time-lapse imaging of the same animal over 12 hr shows statistically insignificant photobleaching of mitochondrial fluorescence ($p$-value 0.055, n = 4 animals, 5 mitochondria, and a box drawn close to each mitochondrion was used to calculate the statistics). Data represented as mean ± SD.

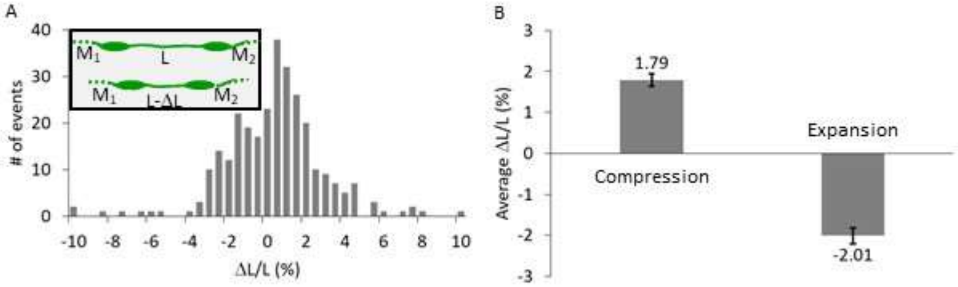


**Supplementary Figure 3:** Effect of repeated immobilization on intermitochondrial distances between stationary mitochondria present on the TRNs. ***A***, Number of events for the percentage of compression or expansion measured from relative intermitochondrial distances ($\Delta L/L$, *n* = 292 total number of events). The inset shows a pair of stationary mitochondria ($M_1$ and $M_2$) with an intermitochondrial distance of L compressed to L-$\Delta$L. ***B***, Average compression ($M_1$ and $M_2$ appear closer) or expansion ($M_1$ and $M_2$ move further apart) percentage values. The data represented as mean ± SEM (*n* = 8 animals imaged for 3 successive time points at 5 min intervals).

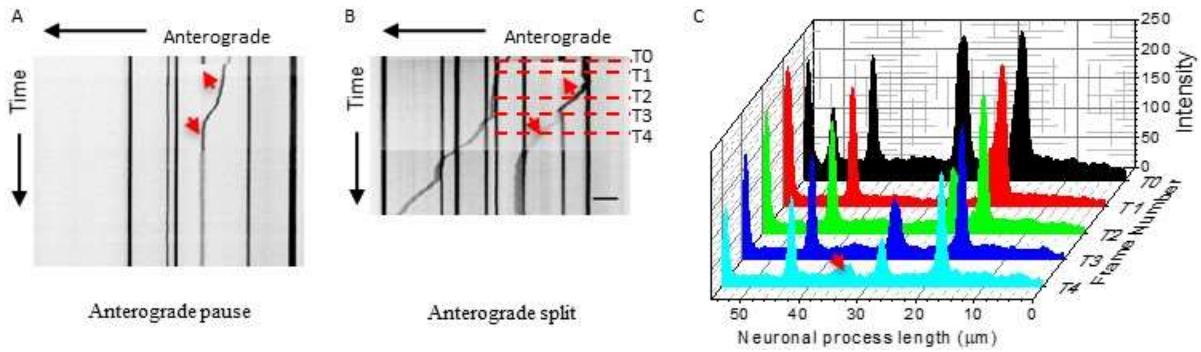


**Supplementary Figure 4:** Kymographs of L4 stage *C. elegans* showing anterograde moving mitochondria across bleached mitochondria. ***A-B***, Anterograde moving mitochondria across bleached mitochondria represented by a red up arrow that pauses (***A***) and partially splits (***B***) at the site of the bleach and represented by red down arrows. The scale bar is 10 μm. ***C***, The intensity distributions along the neuronal process length and at five different time points ($T_0$, $T_1$, $T_2$, $T_3$, and $T_4$), represented by the red dotted lines on (***B***).

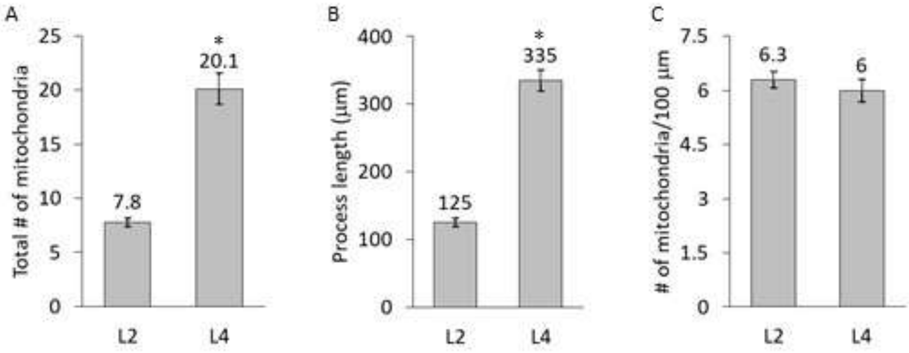


**Supplementary Figure 5:** Mitochondrial density in L2 and L4 animals measured from microfluidic device immobilization. ***A***, Total number of mitochondria counted from L2 (n = 15) and L4 (n = 10) worms immobilized in microfluidic devices. ***B***, Total neuronal process length quantified from the soluble GFP signal. ***C***, Number of mitochondria per 100 μm neuronal process length. The data represented as mean ± SEM. The *p*-value < 0.001 (*) is calculated using two-tailed student t-test. The density of mitochondria in L2 and L4 animals are statistically insignificant.

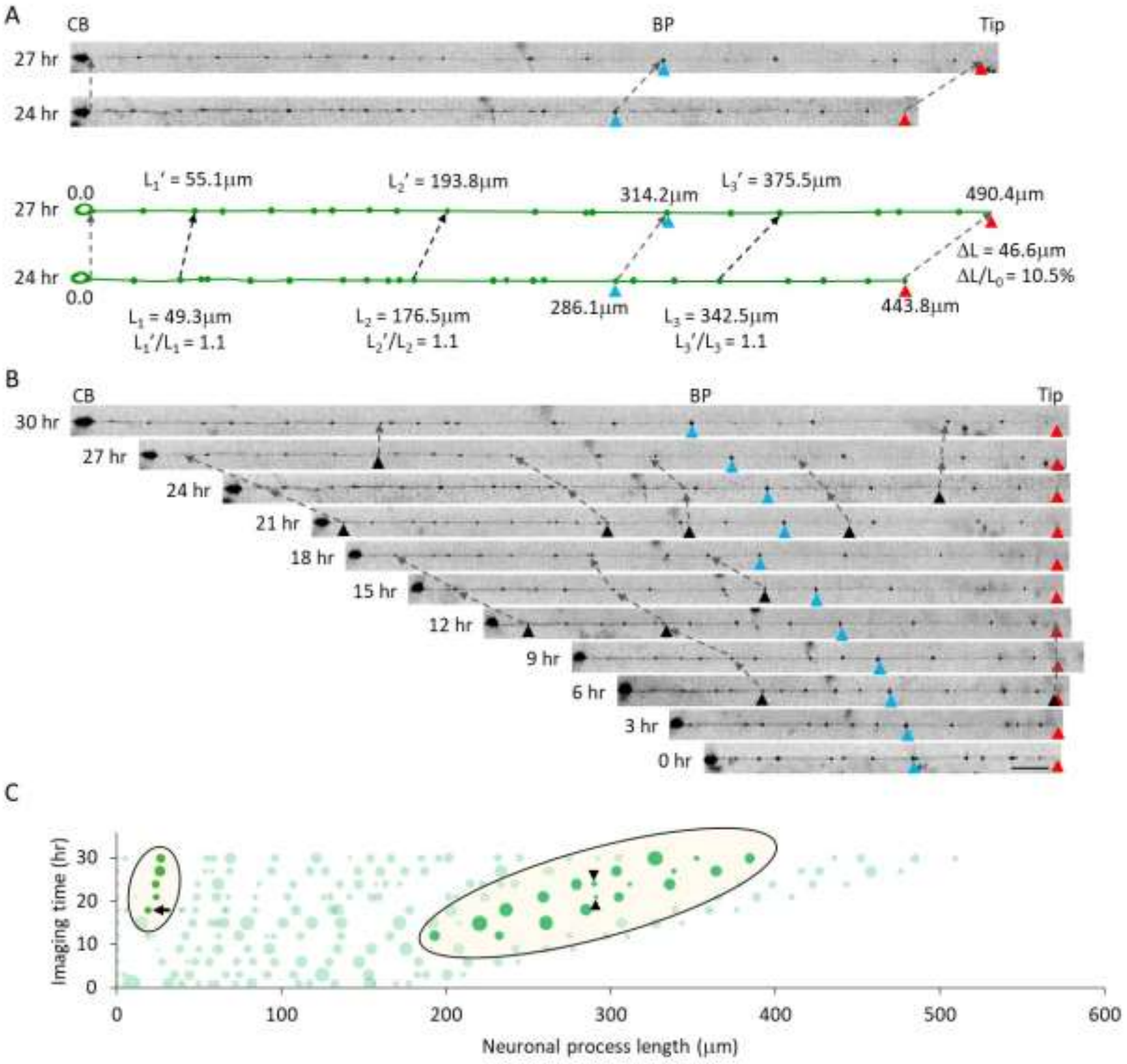


**Supplementary Figure 6:** Time-lapse images of the neuron from an individual animal. ***A***, The images of the entire neuronal process are shown at 24 and 27 hr time points. Two images are aligned with respect to the cell body (CB) and the two additional reference points- the branch point (BP, blue triangle) and tip (red triangle) are marked on the images. The CB, BP, Tip are connected with gray dotted arrows. The trace below the images represents the neuronal processes for both time points with the mitochondria indicated as green dots on respective neuronal processes. The position of the fiduciary markers (CB, BP, and Tip) and three mitochondria are indicated at time t =24 hr ($L_1$, $L_2$, and $L_3$) and t =27 hr ($L_1'$, $L_2'$, and $L_3'$) and connected using gray dotted arrows. The ratio of the two distance between given mitochondria from the cell body at 24 hr and 27 hr corresponds to the observed 10% (46.6 μm) increase in total neuronal process growth ($\Delta L$ = 46.6 μm) in this 3 hr period over the initial process length of $L_0$ = 443.8 μm at t = 24 hr. ***B***, Alignment of the neuronal process image of figure 4(f) for all 11 time points using the tip of the neuron as a reference. ***C***, A representative bubble plot of all 11 time points with the diameter of the bubble proportionate to the total area of each mitochondria. The highlighted region shows the addition of a new smaller mitochondrion (black arrow and black triangles) between two identified mitochondria. The small mitochondrion added on $18^{th}$–time point (black arrow) increases in size in successive imaging frames. A small mitochondrion added at the $9^{th}$-time point (black inverted triangle) did not persist over 6 hr and was not considered as a real event of mitochondria addition.

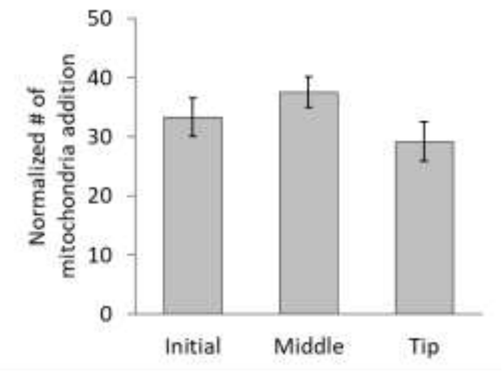


**Supplementary Figure 7:** Normalized number of mitochondria addition events at the initial, middle, and tip of the neuronal process. The normalized number of mitochondria added at each segment of the neuron where the segment corresponds to an equal third of the neuronal process. The total number of mitochondria addition events are 85 (9, 10, 10, 10, 11, 15, 14, and 6 events for each of the eight animals imaged). Data represented as mean ± SEM. The $p$-value $\geq 0.7$ estimated from Student's t-test for each pair indicates that the mitochondria are added uniformly along the neuronal process length.

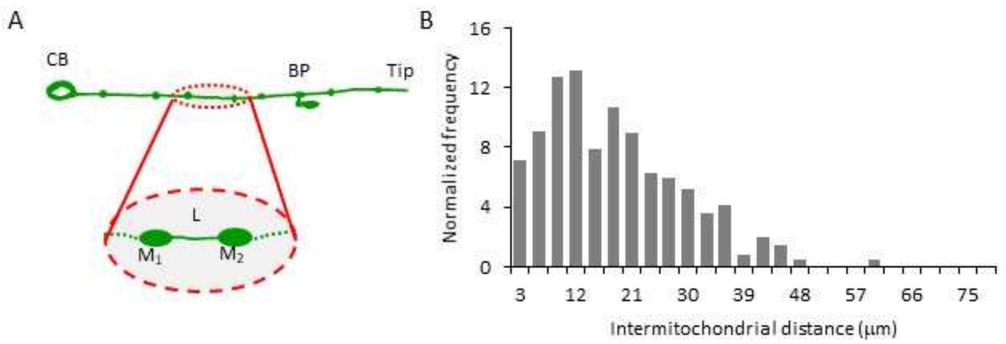


**Supplementary Figure 8:** Intermitochondrial distance measured from L4 stage animals immobilized in the microfluidic chip. ***A***, The schematic of the mechanosensory neuron with cell body (CB), branch point (BP), and tip. The inset shows one of the pair of mitochondria ($M_1$ and $M_2$) with an intermitochondrial distance L. ***B***, Normalized number of intermitochondrial distances in L4 stage animals (n = 8 animals).

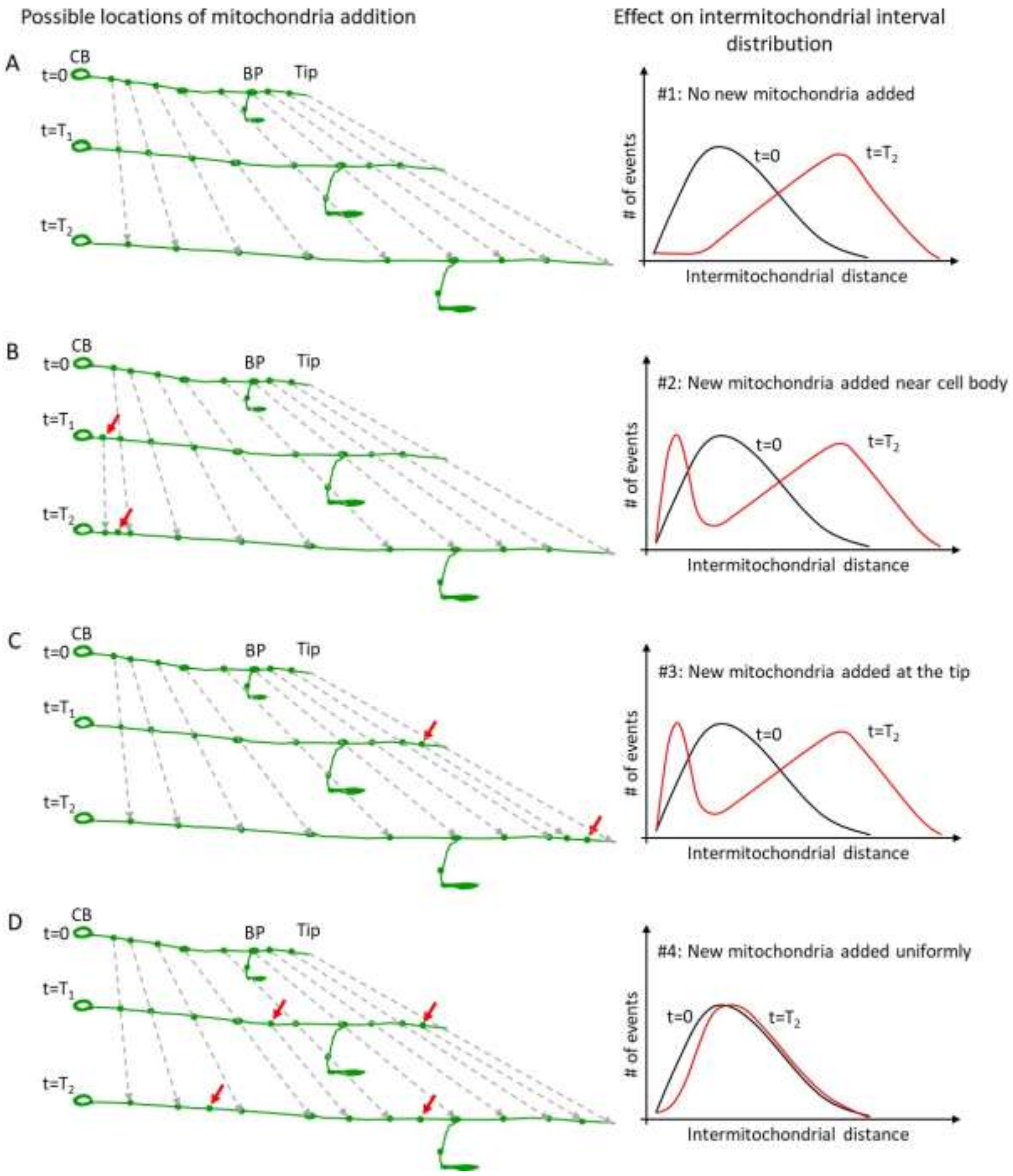


**Supplementary Figure 9:** Schematic representation for mitochondria addition events between pre-existing mitochondria in a growing neuronal process and corresponding changes in inter-mitochondrial intervals. No new mitochondria added (***A***), new mitochondria added near the cell body (***B***), new mitochondria added at the end of the neuron (***C***), and new mitochondria added uniformly along the neuronal process (***D***), and occurs when the intermitochondrial distance between adjacent mitochondria increases beyond a threshold. Three different time points are denoted (t=0, t=$T_1$, and t=$T_2$). The red arrow points to newly added mitochondria. The dashed arrows are added to highlight the position of each stationary mitochondria in successive time points. The panel on the right is a schematic representation of the expected change in intermitochondrial distances for each possible addition scheme.

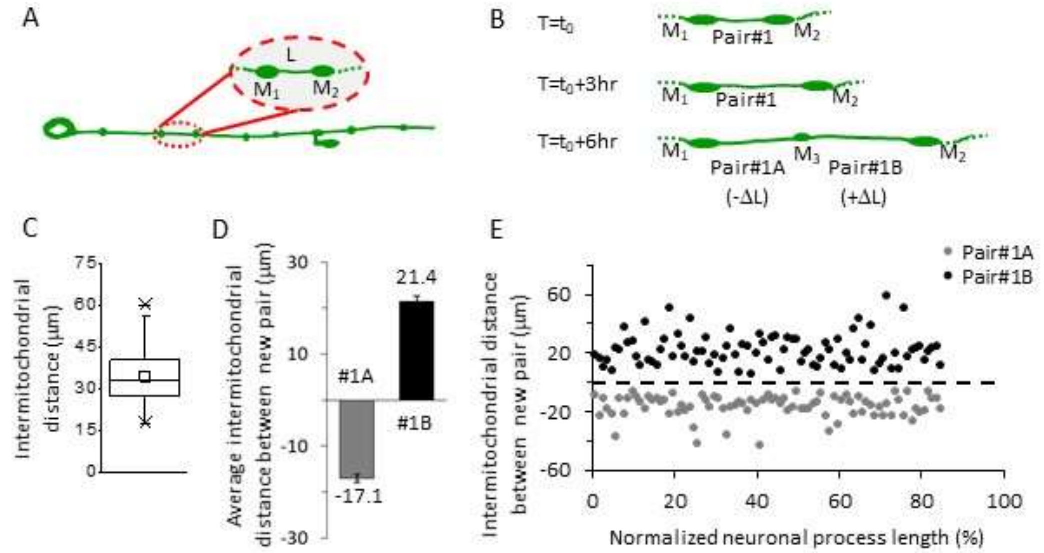


**Supplementary Figure 10:** Intermitochondrial distance statistics when new mitochondria are added in between a pair of old mitochondria. ***A***, Schematic of a mechanosensory neuron with the inset highlighting a mitochondria pair along the neuronal process. ***B***, Schematic of the neuronal process showing the increase in the intermitochondrial distance between the pair of mitochondria at three different time points ($t_0$, $t_0$ + 3hr, and $t_0$ + 6hr). A new mitochondrion is added at $t_0$ + 6hr. The intramitochondrial distance is measured between the old (Pair#1) and new pairs (Pair#1A and 1B) of mitochondria at time points before and after the addition of the new mitochondria, respectively. ***C***, The box and whisker plot all the intermitochondrial distances from time-lapse imaging of the neuronal process imaged from n = 8 animals growing inside the microfluidic chip and imaged for 36 hr. The box represents 25% and 75%, whiskers represent outliers, and the center small box represents the mean (n = 85 mitochondria pair was considered). ***D***, The bar chart of intermitochondrial distances between the new pairs of mitochondria after the addition of a new mitochondrion between a pair of old mitochondria (n = 85 new pairs). ***E***, The intermitochondrial distance for all 85 new pairs (Pair#1A and 1B) are plotted as a function of the normalized neuronal process location, where the addition event occurred.

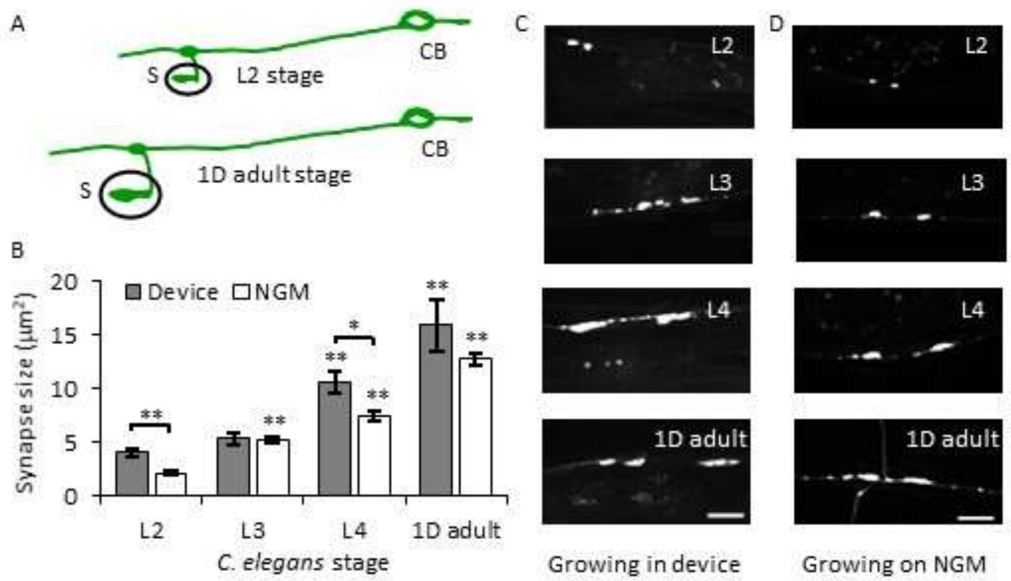


**Supplementary Figure 11:** High-resolution imaging of GFP::RAB-3 in a developing *C. elegans* inside the microfluidic device. ***A***, Schematic of PLM neuron with its cell body (CB) connected to its synapse (S) through a neuronal process at L2 and 1D adult stage of development. ***B***, The plot of synapse sizes at different developmental stages of *C. elegans* growing inside a microfluidic device and on NGM plate. Data represented as mean ± SEM. Statistics are calculated using a two-tailed t-test showing p-value<0.05 (*) and p-value<0.005 (**). The number of synapses analyzed in NGM is n ≥ 37 and in devices are n = 18 (L2), 16 (L3), 14 (L4), and 8 (1D adult). ***C-D***, Images of PLM synapses at L2, L3, L4, and 1D adult stages of development growing inside a microfluidic device (***C***) and on NGM plate and imaged on agar pads (***D***). The scale bar is 10 μm.

**Supplementary Table 1: Turnover of mitochondria events at stationary mitochondria in the PLM neurons of L4 animals immobilized with 3 mM levamisole.**

| | Proximal PLM | Mid PLM |
|---|---|---|
| Anterogradely moving cross | 36% | 43% |
| Retrogradely moving cross | 17% | 18% |
| Anterogradely moving contributes to recovery | 26% | 18% |
| Retrogradely moving contributes to recovery | 21% | 21% |
| Total number of mitochondria analyzed | 88 | 109 |
| Total movies | 29 | 37 |
